# Parsing GTF and FASTA files using the eccLib Library

**DOI:** 10.1101/2025.04.24.650481

**Authors:** Tomasz Chady, Zuzanna Karolina Filutowska

**Affiliations:** Faculty of Mathematics and Computer Science, Adam Mickiewicz University, ul, Uniwersytetu Poznańskiego 4, 61-614, Poznań, Poland; Faculty of Biology, Adam Mickiewicz University, ul, Uniwersytetu Poznańskiego 6, 61-614, Poznań, Poland

## Abstract

**Summary:** Leveraging the Python/C API, eccLib was developed as a high-performance library designed for parsing genomic files and analysing genomic contexts. To the best of the authors’ knowledge, it is the fastest Python-based solution available. With eccLib, users can efficiently parse GTF/GFFv3 and FASTA files and utilise the provided methods for additional analysis.

**Availability and Implementation:** This library is implemented in C and distributed under the GPL-3.0 licence. It is compatible with any system that has the Python interpreter (CPython) installed. The use of C enables numerous optimisations at both the implementation and algorithmic levels, which are either unachievable or impractical in Python.

**Supplementary information:** This library is available for installation from the Python Package Index (PyPI) under the name eccLib https://pypi.org/project/eccLib/. The source code is available at https://gitlab.platinum.edu.pl/eccdna/eccLib. More detailed documentation can be accessed at https://gitlab-pages.platinum.edu.pl/eccdna/eccLib/.

## Introduction

GTF/GFF version 2 and FASTA file formats are widely used in genomics owing to their simplicity and ease of use. Consequently, parsing these files is often the first step in many projects, and can become a bottleneck. Indeed, this was the experience of the authors, who found a Python-based parser to be too slow and memory-intensive for their requirements. To address this issue, numerous libraries exist to facilitate this process. The `gffutils` module (Dale, 2011) parses GTF files into an SQLite database. The `gtfparse` module (Openvax, 2015), the Biopython-affiliated `bcbio-gff` module (chapmanb, 2013), and the GTF parser within the PyRanges library (Stovner and Sætrom, 2019) all return representations of GTF files. For FASTA files, the Biopython library includes a parser called `SeqIO`, and the standalone library `fastaparser` (Kronopt, 2020) is also capable of parsing FASTA files.

However, the performance of some of these libraries may be suboptimal. To overcome these limitations, `eccLib`, a performance-oriented library written in C for Python was developed. The primary objective of `eccLib` is to accelerate the parsing of GTF and FASTA files in Python while providing efficient data structures for genomic data. Users can expect significant performance improvements, particularly with large datasets.

## Methods

The standard implementation of the Python interpreter, CPython, provides an interface for integrating C code, known as the Python/C API. Using this API, developers can write C code that, once compiled and linked to the Python interpreter, can be invoked directly from Python code. This is exactly the interface utilised by `numpy` (Harris et al., 2020) to deliver excellent performance for numerical computations. Development of `eccLib` was not without its challenges owing to its low-level nature. Debugging was particularly problematic when C memory errors propagated into Python reference counting errors. Despite this, the performance gains – even from a direct port of existing Python code into C – can be significant.

However, the true performance improvements arise from low-level access to memory, which permits optimisations that are either unachievable or impractical in Python. Within `eccLib`, this low-level access was utilised to optimise the parsing of GTF and FASTA files, primarily by reducing the number of memory allocations to a minimum and avoiding unnecessary data copying, which is a common issue in high-level languages such as Python.

### Data Types

In order to gain access to the internals of data structures so that they may be optimised, multiple custom data types were created.

The `GtfDict` class is a `Mapping` object used to store data from GTF files. The core GTF entry fields are stored as separate members within the object, permitting access without a hash map lookup. For storing GTF attributes, a custom C hash map is employed, which is faster than a Python `dict`. This hash map, from the `hashmap.h` library (Warden, 2010), utilises the `xxHash` hashing function (Collet, 2012), thereby improving overall performance. Additionally, the `GtfDict` class includes several methods for basic data analysis.

The `FastaBuff` class is used to store FASTA DNA data. The set *I* of IUPAC nucleotide codes (Cornish-Bowden, 1985), can be mapped to a power set of DNA bases using *f* : *I →* 𝒫 ({*A, C, T, G*}). This power set can then be mapped to a 4-bit integer, where each bit represents the presence of a specific base, using *g* : 𝒫 ({*A, C, T, G*}) *→* {0, 1}^4^. The composition of these two functions allows for the efficient storage of sequences as compact binary data, specifically as unsigned 4-bit integers. In this implementation, these 4-bit integers are packed into bytes, enabling a memory usage reduction of at least 50% compared to Python `str` objects, with minimal performance overhead. For instance, the sequence “CG” may be represented as (01000010)_2_, and the sequence “DNT” as (10111111 00010000)_2_. For non-nucleotide characters, the Python `str` object is used as a fallback.

### GTF parsing

The principal challenge in GTF parsing lies in string tokenisation and the optimal storage of attributes. To address this, a custom C tokeniser that does not modify the input data is utilised – thereby avoiding additional memory allocations – and a third-party hash map library is employed for caching and attribute storage. Caches are utilised for storing already encountered keys and values during parsing, thus avoiding redundant memory allocations for identical strings. In addition, the process of resolving URL and UTF-8 encodings was optimised by using a custom C function that simultaneously resolves both encodings, which significantly increases overall performance.

### FASTA parsing

Parsing the FASTA format is considerably simpler than GTF processing. The primary challenge concerns the memory storage of sequences. To address this, the `FastaBuff` class is employed as the storage data type by the primary FASTA parser, `parseFASTA()`. This approach reduces the memory usage of the FASTA parser compared with implementations that use Python `bytes` or `str` objects.

## Results

To assess the performance of the `eccLib` library, a benchmark was conducted. This benchmark was performed on a Linux system equipped with an Intel i9-9900KF CPU, 48.0 GB of RAM, and an SSD. The objective was to measure how quickly each parser could return a representation of the provided file. For iterative parsers, the outputs were appended to a list, thereby simulating the behaviour of a non-iterative parser. In turn, 20 new Python instances were launched for each parser, measuring the time with the `datetime` module and memory usage with the GNU Time utility. To better illustrate overall performance, the Time-Memory Product (TMP) was calculated using the formula TMP = Time × Memory, in appropriate units.

### GTF parsing benchmark

Benchmark results suggest that the GTF parsing methods in `eccLib` significantly outperform both `gtfparse` and the parser within the PyRanges library. The benchmark file used for speed tests was the 1.4 GB `Homo sapiens.GRCh38.108.gtf` from Ensembl (Dyer et al., 2024), chosen for its size and accessibility. Parsing methods within `eccLib` were compared against one another, the `gtfparse read gtf` function, and the PyRanges `read gtf` function. The results are presented in fig. 1.

**Fig. 1.**
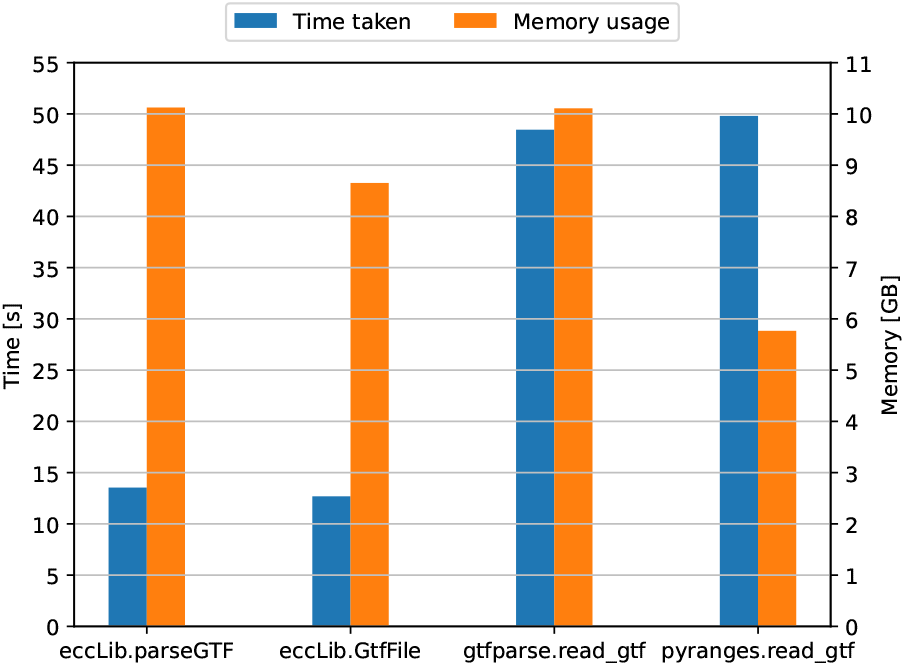
Speed and peak memory usage of different GTF parsing methods. The methods presented here, `parseGTF` and `GtfFile`, achieved TMP scores of 137 and 110, respectively. Third-party methods, `gtfparse.read_gtf` and `pyranges.read_gtf`, achieved TMP scores of 490 and 287, respectively.

The results indicate that methods presented here are significantly faster at parsing GTF files compared to the alternatives, albeit with higher memory usage. `eccLib.GtfFile` parser is 3.9 times faster than `pyranges.read_gtf`, while consuming 1.5 times more memory.

Contrary to expectations, the iterative parser was not significantly slower than the non-iterative parser. The authors attribute this to the hardware characteristics, which mitigated the performance penalty incurred by additional I/O operations.

One factor that may have influenced the results is the high number of repeated GTF attribute keys and values in the benchmark file; however, the authors do not believe this had a significant impact on the overall outcomes. Unfortunately, `gffutils` and `bcbio-gff` could not be tested due to their poor performance on the test file.

### FASTA benchmark

Testing indicates that the `parseFASTA()` function in `eccLib` outperforms contemporary FASTA parsers. The benchmarking was conducted using the same methodology as the GTF parsing benchmark. The test file used was the 3.2 GB `Homo sapiens.GRCh38.dna.primary assembly.fa` from Ensembl. The averaged results are shown in fig. 2.

**Fig. 2.**
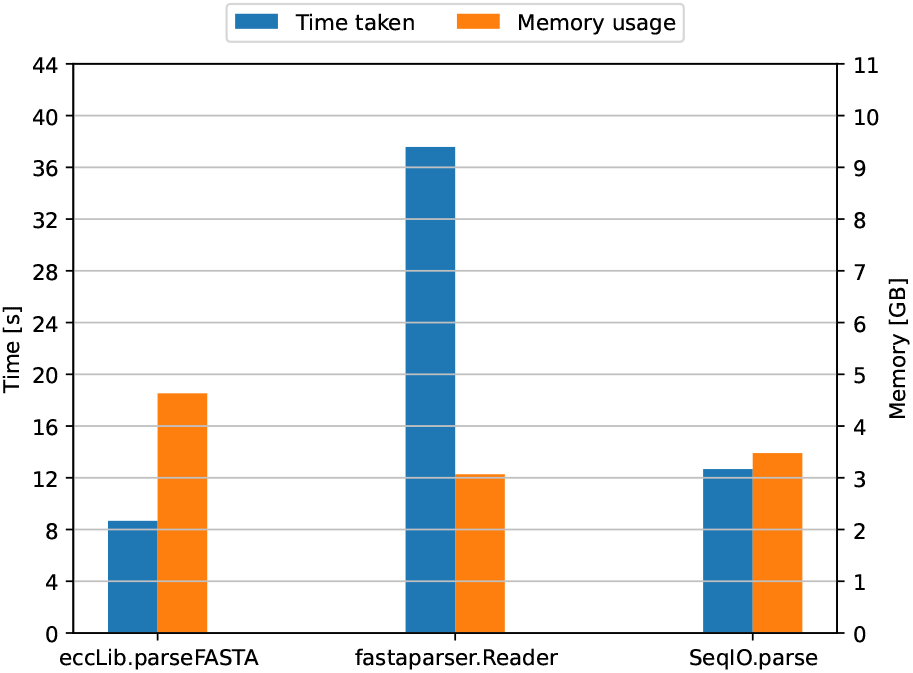
Speed and peak memory usage of different genomic FASTA parsing methods. The function presented here, `parseFASTA()`, achieved a TMP of 40, while `fastaparser` and`SeqIO` achieved TMP scores of 115 and 44, respectively.

While `parseFASTA()` demonstrates a clear speed advantage, it incurs increased memory consumption. The `fastaparser` reader was also used in “quick” mode to manage memory usage.

Since `eccLib` handles DNA sequences differently from protein sequences, a separate benchmark was conducted using the 85.1 MB `Homo_sapiens.GRCh38.pep.all.fa` from Ensembl. The results are illustrated in fig. 3.

**Fig. 3.**
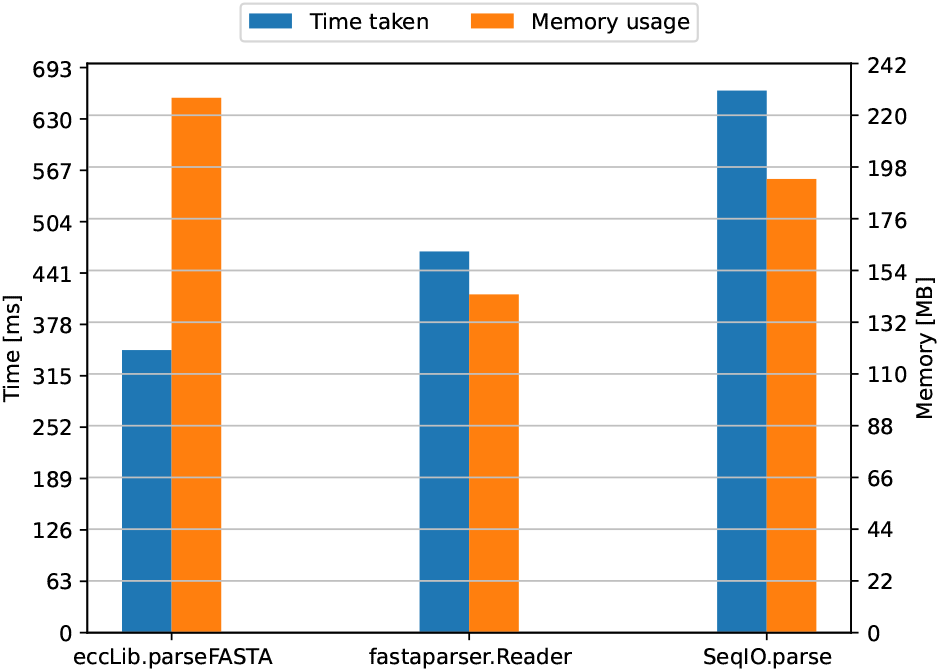
Speed and peak memory usage of different protein FASTA parsing methods. `fastaparser.Reader` achieved the best TMP of 67, with eccLib and SeqIO achieving TMP scores of 79 and 128, respectively.

Here, the relative performance of `eccLib` is notably poorer; however, the speed advantage remains; the increased memory usage is largely attributable to the fact that protein sequences are not stored as efficiently as genomic sequences.

## Discussion

The `eccLib` library provides a high-performance solution for parsing GTF and FASTA files in Python, thereby significantly enhancing project performance. Although it utilises more memory than alternative approaches, this is a consequence of its adoption of hash maps rather than fixed-field tables, which better accommodate the flexibility of GTF/GFF files. Despite the higher memory usage, the performance benefits are substantial, and the authors contend that these advantages justify the increased memory consumption. Future work will focus on expanding features and improving interoperability with other libraries. The choice of C as the implementation language was driven by the need for performance. However, the authors acknowledge the inherent issues associated with programming in C. The authors find the possibility of developing a library for Python with a similar feature set in a higher-level language, such as Rust, to be an interesting avenue for future exploration.

## Acknowledgements

This work was supported by a grant from the Polish Ministry of Education and Science (MEIN) [0054/DIA/2014/43].

The authors declare that they have no competing interests.

No new data were generated or analysed in support of this research. Data used for benchmarking were obtained from Ensembl (Dyer et al., 2024).

